# Nitrogen dynamics and yield performance of an elite bread wheat line with BNI capacity expressed in an alkaline soil

**DOI:** 10.1101/2025.07.29.667244

**Authors:** Hannes Karwat, Masahiro Kishii, María Elena Cárdenas-Castañeda, Maria Itria Ibba, Victor Kommerell, Alison R Bentley, Hans-Joachim Braun, Iván Ortiz-Monasterio

**Affiliations:** International Maize and Wheat Improvement Center (CIMMYT), Mexico-Veracruz, El Batán, Texcoco CP 56237, Estado de Mexico, Mexico; Japan International Research Center for Agricultural Sciences (JIRCAS), 1-1 Ohwashi, Tsukuba, Ibaraki 305-8686, Japan; Research School of Biology, Australian National University: Canberra, Australia

## Abstract

Wheat with biological nitrification inhibition (BNI) has been shown to be effective in acidic soils. BNI expressions in alkaline soil have not been documented in field studies. We present the first observation of BNI effects in alkaline soil trials (pH 8.6-8.7) of elite spring wheat that carries the Lr#N short arm from *Leymus racemosus*.

BNI was expressed as lower soil nitrate in three trials conducted in an irrigated wheat system in northwestern Mexico. In one year, we observed 24-37% lower nitrate in soil of the translocation line compared to the control after the second ammonium-N fertilizer application. Lower nitrate was observed in the row and in the furrow. The largest difference between the control and the translocation line was approximately one month after the second ammonium-N split application (73-77% less nitrate). Potential nitrification rates were 27-32% lower in soil from the translocation line compared to the control, one week after a high ammonium-N application.

Higher flag leaf nitrate concentration of the translocation line could be related to the strongly reduced soil nitrate. The translocation line took longer to reach anthesis and flowering. In two experiments, the translocation line was equivalent in terms of grain yield, whereas in one experiment it produced lower yields, with fewer grains per spike and reduced number of spikes per area. Protein content and grain N uptake of the translocation line were similar to the control. We conclude that BNI reduces nitrification with a spatially and temporally significant impact under alkaline soil and high N input conditions. This finding points to the potential major environmental benefits that could be achieved also in non-acidic soil spring wheat systems worldwide. The effect of the Lr#N short arm on yield quantity and quality in other elite lines needs further investigation.

## Introduction

Global wheat production consumes about 20% of total applied nitrogen (N) fertilizer (Ladha et al. 2016). The calculated average N use efficiency (NUE = [plant N removed) - (soil mineral N + N deposited in rainfall)] / (fertilizer N applied)) of cereal systems was 33% two decades ago (Raun and Johnson 1999). Twenty years later, a new assessment showed that NUE did not increase significantly on a global scale (Omara et al. 2019): 67% of the applied N fertilizer does not contribute to wheat yield formation but is either bound in the soil profile or released in reactive forms, through leaching and/or gaseous forms to the environment. The microbial conversion of ammonium (NH_4_ ^+^) to nitrate (NO_3_ ^-^), is associated with high N losses via NO_3_ ^-^ leaching. Furthermore, nitrate is the substrate for denitrifiers. Both processes are responsible for the emission of N_2_O directly from fields and downstream water bodies with considerable contributions to global warming (Nevison, 2000). Improved agronomic N management in wheat systems includes using right formulation, rate, application time and place and can reduce N losses already tremendously, but their scaling out has proved challenging both in The Global North and South. BNI is a seed-based intervention and could expand the options of agronomic interventions to optimize NUE. In low- and middle-income countries (LMIC), scalable Ag-based mitigation solutions are often seed-based. Consequently, breeding for low N requirement genotypes must go hand-in-hand with context-specific, optimized N application techniques and strategies (Swarbreck et al. 2019).

Wheat landraces and wild relatives, as donors for resistance to biotic and abiotic stress, also constitute a wheat breeding pool with high potential for improving plant N fertilizer uptake (King et al. 2024) and reducing N losses to the environment. By harnessing soil-root interactions (Baggs et al. 2023) and translating the known nature-based solution into breeding approaches (Reynolds et al. 2021), N footprints of wheat systems can potentially be improved. Coping mechanisms developed by wild relatives to survive in N scarce environments, like biological nitrification inhibition (BNI), may alter both the plant N uptake and N preservation of the plant-soil-microbe system (Lata et al. 2022). Traits that are responsible for the production and release of nitrification inhibitors to the soil are located on the short-arm of ‘Lr#N’ chromosome (Lr#N-SA) of crop wild relative *Leymus racemosus* L (Subbarao et al. 2021). By introducing this trait into high yielding bread wheat lines, they benefit from an extended period of ammonium availability due to BNI and a more balanced ammonium-to-nitrate nutrition (Subbarao and Searchinger 2021). The slower below-ground conversion of ammonium to the very mobile form of nitrate and lowering ammonia oxidation will reduce the release of the most crucial greenhouse gas in Agriculture, N_2_O (Prosser et al. (2020).

BNI release from roots in hydroponics are triggered by NH_4_ ^+^; this increases significantly under lower pH conditions (Subbarao et at. 2007; Egenolf et al. 2021). Proof-of-concept field and pot studies reported that the BNI trait introduction 1) translated into higher grain yields, 2) improved plant N uptake, 3) reduced nitrate formation and 4) significantly lowered N_2_O emissions, based on soil samples where BNI-wheat was sown (Subbarao et al. 2021, Bozal-Leorri et al. 2022). The most promising field results derived from experiments with a soil pH 5.0 to 5.5 (Subbarao et al. 2021) and pot studies that used soil with a neutral pH of 7 (Bozal-Leorri et al. 2022). However, more than half of the soils devoted to wheat cultivation in the Global South are alkaline.

The Yaqui Valley in Mexico, with soil pH of 7 to 8.5, should be an excellent testing ground for BNI expression in a high N input spring wheat system with alkaline soils (Ahrens et al. 2012). Nitrate leaching for this spring wheat system in northwestern Mexico occurs within the range of 2 to 5% of applied N per crop season on-research station and 14-26% in farmers’ fields. Farmers typically apply between 250-300 kg N ha^-^ (Riley et al. 2001). BNI has been shown to reduce nitrate leaching by the tropical forage grass *Brachiaria humidicola* (Karwat et al. 2018) but comparable studies for BNI-wheat are wanting.

Ou research aimed to confirm the BNI potential of ROELFS-Lr#N-SA, earlier established with a bioluminescence assay under controlled conditions (Subbarao et al. 2021), in field experiments under high soil pH and N fertilization conditions, by assessing yield quantity and quality stability of the translocation line.

Our hypotheses were, that 1) the BNI potential is expressed as lower nitrate values in the soil of plots sown with ROELFS-Lr#N-SA compared to ROELFS (control); 2) that potential nitrification rates (PNRs) via soil incubation assay would reflect a BNI effect in vitro from soil samples taken after field ammonium fertilization; 3) that soil ammonium levels would be higher due to BNI expression compared to levels in soil of the non-BNI wheat line; We further conjecture that 4) lower soil nitrification is reflected in lower plant tissue nitrate of the translocation line; Furthermore, 5) ammonium addition triggers both the release of BNI by the root and kick-off soil nitrification and illustrates the differences among the translocation and control line; Finally, we hypothesized that 6) Lr#N-SA would neither disrupt yield potential nor end-use quality traits.

## Material and Methods

### Field location

Three field experiments were conducted at the Experimental Station Norman E. Borlaug (CENEB) in the Yaqui Valley of Northwestern Mexico, Ciudad Obregón in the state of Sonora (27◦24’N,109◦5’W, 38m) from 2019 to 2022. The area is Mexico’s most important spring wheat production area resembling climatic (dry arid) and crop management (irrigated high input) conditions of the mega-environment 1 (Rajaram et al. 1993; Braun et al. 2010). Dominant soil types in the Yaqui Valley are Aridisols and Vertisols, low in organic matter (<1%) with a slightly alkaline to alkaline soil pH range (Limon-Ortega et al. 2002). In the Yaqui Valley, spring wheat is sown in November and December and harvested in May, followed by a fallow period. Wheat is sown on beds with irrigated furrows. N fertilizer inputs of 250-300 kg N ha^-1^ applied as urea and N anhydrous ammonia (NH_3_) are common for this region (Ortiz-Monasterio and Raun, 2007; Fischer et al. 2022).

### Sowing material

The elite spring wheat line for the field trials was ROELFS F2007 (ROELFS), a leading bread wheat variety with stripe rust and leaf rust resistance developed for southern Sonora, Northwestern Mexico in the late 2000’s to early 2010’s (Figueroa-Lopez et al. 2008). Chromosome addition lines have been developed as outlined by Kishii et al. (2004), the complete short-arm chromosome (Lr#N-SA = T3BL.3NsbS) transferred as reported by Subbarao et al. (2021) and was backcrossed five times. ROELFS-Lr#N-SA (CSMONO3B/3/CS/LE.RA/CS/4/CSph/5/5*ROELFS(N)) showed a BNI in vitro capacity of 162.2 allylthiourea units (ATU) g^-1^ root dry weight d^-1^ (Subbarao et al. 2006; 2021) compared to 86.5 ATU for ROELFS. The presence of the translocation on chromosome 3B was confirmed in harvested seed samples by genomic in situ hybridization (GISH, as per Kishii et al. 2004). The two genotypes ROELFS (control) and ROELFS-LR#N-SA (BNI-line) were used in the trials.

### Experiment 1 (2019/2020)

Sowing was conducted on 21 December 2019. Each experimental plot contained four beds 0.8m wide and 2.5m long. The two central beds were kept for the harvest area whereas the first and last beds were used for soil and plant tissue sampling. The two genotypes ROELFS (control) and ROELFS-LR#N-SA (BNI-line) were sown with a sowing density of 250 seeds m^2^.

The trial was set up as a split-plot design with N rate as main plot and genotype as subplot as the two treatments replicated three times. The experiment was irrigated five times during the crop cycle when the soil water availability reached 50%. Weed control was a combination of mechanical and chemical control. Diseases and insects were controlled by application of respective spray solutions. Days to 50% anthesis and days to physiological maturity were recorded.

Soil from the upper horizon (0-15 cm) had a pH of 8.7. Fertilizer application at sowing included phosphorus (46 kg P ha^-1^ of triple super phosphate (P_2_O_5_) and nitrogen applied to the furrow in the form of ammonium sulphate (NH_4_)_2_SO_4_, 83 kg N ha^-1^. The ‘LOW N’ treatment plots did not receive further N fertilizer whereas the ‘HIGH N’ treatment plots were fertilized again at booting (60 days after sowing). The ammonium sulphate fertilizer was broadcasted by hand on the top of the beds at a rate of 167 kg N ha^-1^.

### Leaf sampling for nitrate tissue determination (Exp 1)

Leaf samples were collected one time before and four times after the 2^nd^ N application in Exp 1. Half of the flag leaf was cut off with scissors and stored in aluminum foil envelopes. Ten leaves were sampled from the two outside beds of each plot. The envelopes were dropped into an insulated liquid N container. and then stored in a freezer at -80 degrees Celsius. All leaves were cut with scissors into 2mm thin stripes and loaded into a garlic press. The NO_3_ ^-^ concentration was determined by optical evaluation of the color scale of Merckoquant nitrate test strips measured with a Nitrachek 404 Meter (Hach).

### Soil sampling, mineral N determination (Exp 1)

According to current understanding (and observed for other BNI plants as well) ammonium addition to the root system increases the release of BNI substances. Furthermore, nitrification peaks during the crop season after fertilization of ammonium-based fertilizers or urea.

Soil samples were collected from the topsoil (0-15cm) horizon between the 2 rows and between the beds (furrow) in 2020 (Exp 1) at 14 and 33 days after the second N fertilizer application. Ten subsamples were taken, mixed in a bucket and sent in a cooled box as representative samples of approximately 250g to the laboratory. NO_3_ ^-^ determination was measured in a yellow-colored aqueous complex, developed by the reaction of sulfuric acid and sodium salicylate in the presence of ammonium sulfamate (to avoid nitrite ion interference). The determination of NH_4_ was based on the formation of a greenish-colored complex, which developed from the reaction between ammonia, sodium hypochlorite, EDTA, and salicylate in the presence of sodium nitroprusside. Absorbance was measured with a Synergy HT spectrophotometer (BioTek Instruments) at 410 nm (NO_3_ ^-^) and at 667 nm (NH).

### Experiment 2 (2020/2021) and Experiment 3 (2021/2022)

Since the N fertilizer effect was not significant in the last season, we moved to plots with lower N for two consecutive seasons. To conduct a more frequent soil sampling, the 2^nd^ and 3^rd^ experiment was established with larger sowing beds of 5 m length. As described for Exp 1, four beds of 0.80m width, sown with two rows per bed were one experimental unit. The phosphorus application rate was 69 kg P_2_O_5_ (N-P-K: 0-46-0) at pre-sowing according to the recommendation of the station management. Soil pH in 0-15cm layer was 8.6. The first N application was conducted as band between the beds, followed by irrigation. The second N doses at booting stage was applied directly within the two rows on top of the beds, also followed by irrigating the furrow. Similar to Exp. 1 we measured soil nitrate and ammonium frequently after the second ammonium-N fertilizer application and after harvest, but only within the row, where plants grew. Additionally, soil samples were collected after N fertilization three times every 7 days and once after harvest and incubated (description below). Exp 2 was sown on 21 November 2020 and Exp 3 on 19 November 2021.

### Soil sampling, mineral N determination (Exp 2 and Exp 3)

In 2021 (Exp 2), soil samples for nitrate and ammonium determination were collected directly from the sowing row at 7, 14 and 21 days after the second N fertilization and at harvest.

For both Exp 2 and Exp 3, soil sub-samples were incubated in triplicates according to a modified protocol of the shaken soil-slurry method by Hart et al. (1994). It is a method that has been successfully used in other BNI research studies (O’Sullivan et al. 2016). Briefly, 10g of fresh soil was added into 200ml volume Erlenmeyer flasks. A volume of 100ml ammonium-sulphate solution with potassium monobasic phosphate (KH_2_P0_4_) was added as N and P source for nitrifiers. The media was adjusted to pH 7.5 as it is considered that nitrification peaks for these conditions. The flasks were placed into an orbital shaker with a controlled incubation temperature of 25°C. Vigorous shaking at 100rpm continuously aerates the slurry and has been shown to prevent denitrification (Hart et al., 1994). Furthermore, due to the low C-to-N rate in the slurry N immobilization is negligible. Consequently, NO_3_ ^-^ consumption in slurry can be excluded and net and gross nitrification rates in this microcosm study do not differ significantly according to Hart et al. (1994).

Sub-samples (5 ml) were taken 1 h and 144 h after incubation initiation, filtered and frozen. NO_3_ ^-^ and NH_4_ ^+^ were measured colorimetric in 96 well-plates as mentioned above. Potential nitrification rates (PNRs) were expressed in N-NO_3_ ^-^ mg^-1^ dry soil g^-1^ day^-1^.

### Estimation of nitrate N amounts in topsoil (Exp 1 and Exp 3)

Topsoil (0-15cm) nitrate in kg N–NO_3_ ^-^ ha^-1^ was calculated based on the measured nitrate concentrations (Fig 1 and Fig 3). The average bulk soil density at CENEB is 1.3 g cm^-3^ (Lobell & Ortiz-Monasterio, 2006). Consequently, the amount of soil in the respective soil horizon was calculated and extrapolated with the nitrate concentration measured and expressed as kg N-NO_3_ ^-^ per ha^-1^. Accordingly, the following equation was used to estimate the range on nitrate-N in the topsoil per hectare (ha):

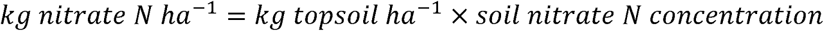

**Fig 1:**
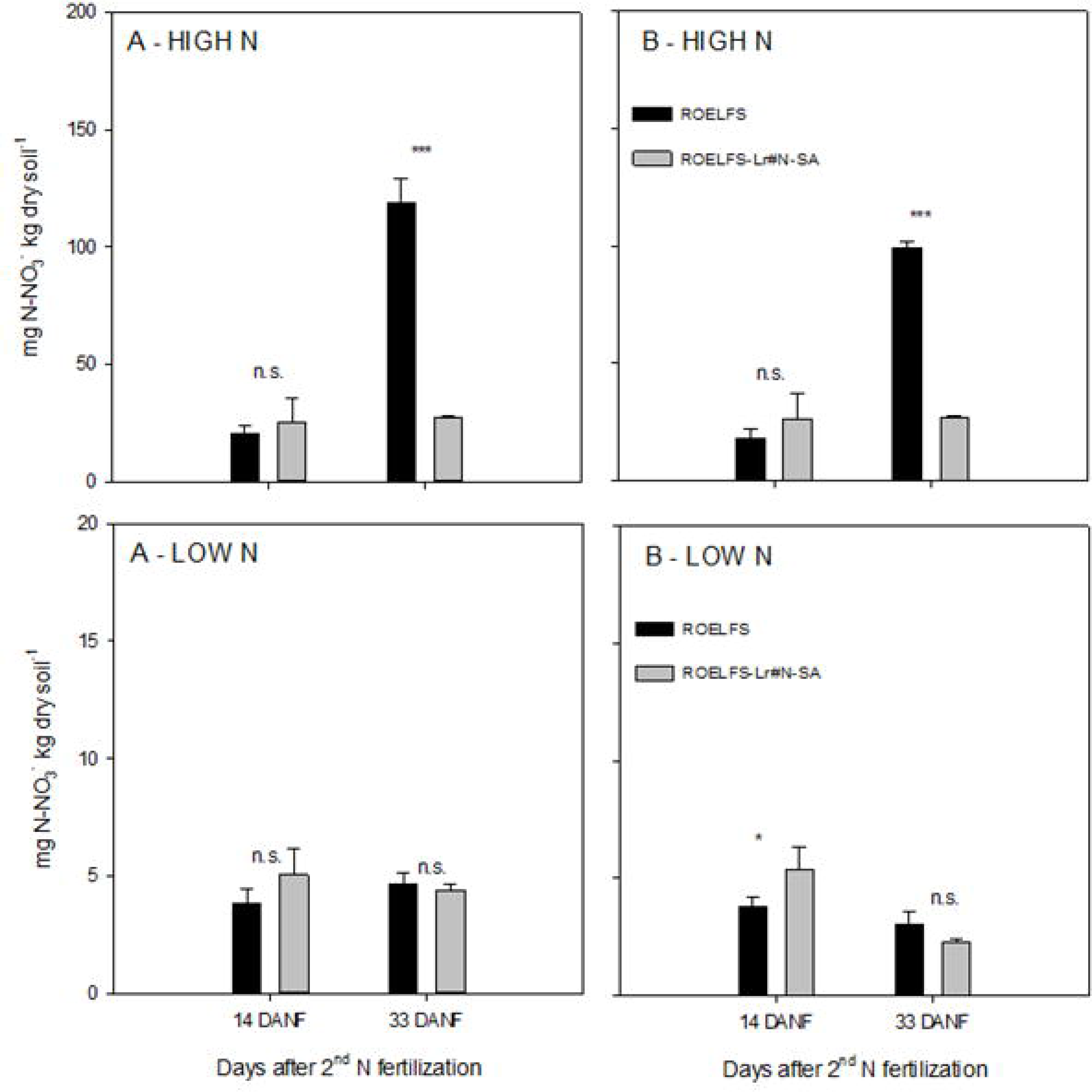
Nitrate (N-NO_3_) concentration in topsoil (0-15cm) in an experimental plot at CIMMYT’s research station in Ciudad Obregon, North Mexico in the field season 2019/2020. HIGH N: ammonium-N fertilizer application was split into 83 kg N ha^-1^ at sowing and 167 kg N ha^-1^ at booting stage. LOW N: ammonium-N fertilizer application as a single application of 83 kg N ha^-1^ at sowing. Samples were taken at 14 and 33 days after the second ammonium fertilizer application (DANF). A = soil samples taken from the sowing beds, between two wheat rows where the N fertilizer has been applied. B = soil samples taken from the furrow (between the sowing beds), where the irrigation water was applied. ROELFS = parental line, control. CSMONO3B/3/CS/LE.RA//CS/4/CS ph/5/5*ROELFS = ROELFS line that carries the Lr#N-SA from *Leymus racemosus*. Error bars display the standard error of the mean of 3 replicated plots. Significant differences for the pairwise comparison for each sampling date separately is indicated for p-levels *<0.01, **<0.001 and ***<0.0001. n.s. = not significant.

Where the amount of soil in the topsoil layer (0-15cm) per ha was calculated according to:

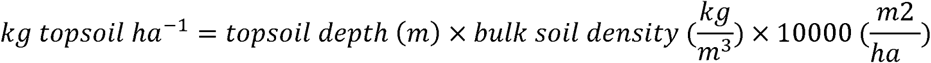

### Wheat quality evaluation (Exp 2 and Exp 3)

Samples from were evaluated for their grain, milling, flour, and end-use quality characteristics. Specifically, to assess the physical and chemical quality of the grain, all samples were analyzed for test weight (TW) according to AACC method 55-10.01 (AACC Approved Methods of Analysis, 2011); thousand kernel weight (TKW) and grain hardness (GRNHRD) using the Single Kernel Characterization System (SKCS) instrument (SKCS Model 4100; Perten Instruments) following AACC method 55-31.01 (AACC Approved Methods of Analysis, 2011); and grain nitrogen (N) content measured using the Dumas combustion method (FP828 Leco® Instruments) following AACC official protocol 46-30.01 (AACC Approved Methods of Analysis, 2011). The total grain protein content was calculated as 5.7 times the total N content (Teller, 1932).

Grain samples were then tempered with water according to the official AACC method 26-95.01 (AACC Approved Methods of Analysis, 2011) and milled into flour using a Brabender Senior experimental mill. Flour protein (%) and moisture content (%) were determined by near-infrared spectroscopy (Antaris FT-NR analyzer, Thermo Fisher Scientific, USA), calibrated as per AACC methods 46-11.02 and 39-11.01, respectively (AACC, 2011).

Overall gluten quality was evaluated through SDS-Sedimentation volume (SDSSED, mL), following the protocol reported by Peña et al., (1990). Gluten strength (alveograph W), and the tenacity/extensibility ratio (alveograph P/L) were determined using 60g flour samples, following the Alveograph manufacturer’s instructions (Chopin, France) and AACC method 54-30.02 (AACC, 2011).

The bread-making process was carried out using the straight dough method with 100 g of flour (AACC method 10-09.01). Bread loaf volume (LV) was determined by rapeseed displacement using a volume-meter. The amounts of water added during alveograph testing, and baking were determined by near-infrared spectroscopy (Antaris FT-NR analyzer, Thermo Fisher Scientific, USA), calibrated according to Guzmán et al. (2015).

Characterization of the glutenin profile was conducted through SDS-PAGE following the methods reported by Guzmán et al., (2022).

### Statistical analysis

All data were analyzed with a mixed model approach with SAS 9.4. With the PROC GLM or PROC GLIMMIX (standard linear mixed model) method. The model considered GTP (genotype), N_RATE (N fertilizer treatment) REP (field or laboratory replication of the observation) and possible interactions. The analysis was conducted for each year and field site separately. Independent variables that were found to have no significant effect (α = 0.05) on the dependent variables were excluded from the mixed model during the pairwise comparisons. Additionally, to compare mean values of the two wheat lines of interest the PROC TTEST was conducted separately for year, field, sampling time and N rate treatment.

## Results

### Soil nitrate and ammonium

In the HIGH N Plots, two weeks after the second NH_4_ ^+^ application (Exp 1) no differences (p>0.05) in terms of soil NO_3_ ^-^ were observed between the two accessions by sampling in the sowing row (Fig 1 A) and in the furrow (Fig 1 B). However, 33 days after the second ammonium sulphate application (DANF), in the ROELFS plots in the sowing row around 119ppm of NO_3_ ^-^ was measured compared to 27ppm NO_3_ ^-^ ROELFS-Lr#N-SA. The data for samples from the furrow (about 40 cm from where the first N rate was applied) were 99 ppm NO_3_ ^-^ for ROELFS vs. 27ppm NO_3_ ^-^ for. ROELFS-Lr#N-SA. Measuring in the LOW N plots showed a NO_3_ ^-^ concentration of 4.3 – 6.6ppm with no differences (p>0.05) among lines, among the sampling dates or when comparing furrow and sowing row. In one case (Fig1 B - LOW N) the translocation line showed significantly higher NO_3_ ^-^nitrate values as the control in the furrow.

Soil NH_4_ ^+^ concentrations in Exp 1 were higher in the sowing rows at 14 DANF compared to 33 DANF (Fig 2A, HIGH N). Significantly lower concentrations (∼4ppm N-NH_4_ ^+^ lower) were measured between the rows at 33 DANF for the translocation line compared to the parental (Fig 2 A, HIGH N). At 14 DANF the soil NH_4_ ^+^ difference between ROELFS-Lr#N-SA and control were ∼20ppm although not statistically significant (Fig 2 B, HIGH N).

**Fig 2:**
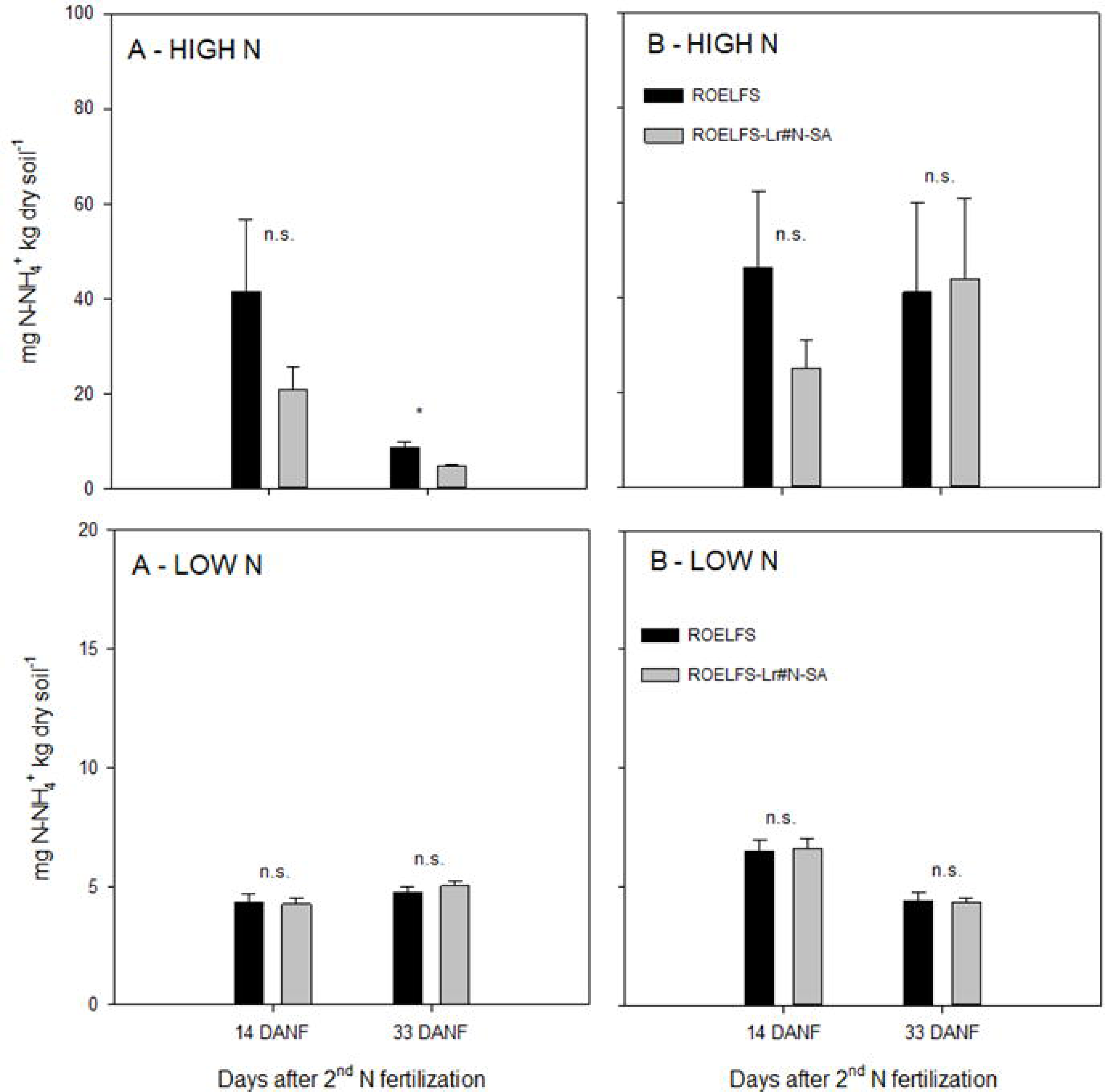
Ammonium (N-NH_4_^+^) concentration in topsoil (0-15cm) in an experimental plot at CIMMYT’s research station in Ciudad Obregon, North Mexico in the field season 2019/2020. HIGH N: ammonium-N fertilizer application was split into 83 kg N ha^-1^ at sowing and 167 kg N ha^-1^ at booting stage. LOW N: ammonium-N fertilizer application as a single application of 83 kg N ha^-1^ at sowing. Samples were taken at 14 and 33 days after the second ammonium fertilizer application (DANF). A = soil samples taken from the sowing beds, between two wheat rows where the N fertilizer has been applied. B = soil samples taken from the furrow (between the sowing beds), where the irrigation water is applied. ROELFS = parental line, control. CSMONO3B/3/CS/LE.RA//CS/4/CS ph/5/5*ROELFS = ROELFS line that carries the Lr#N-SA from *Leymus racemosus*. Error bars display the standard error of the mean of 3 replicated plots. Significant differences for the pairwise comparison for each sampling date separately is indicated for p-levels *<0.01, **<0.001 and ***<0.0001. n.s. = not significant.

**Fig 3:**
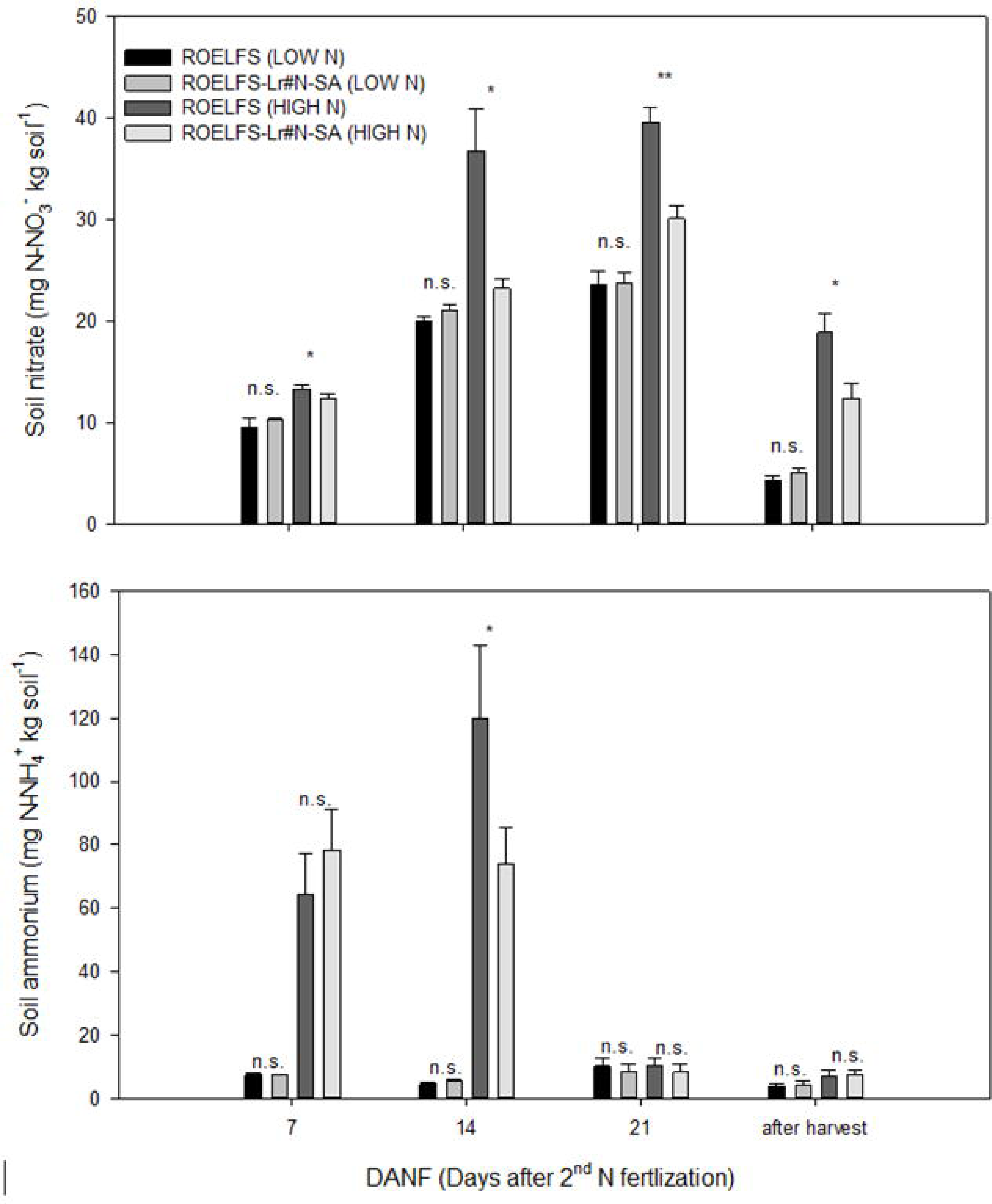
Nitrate (N-NO_3_) concentration in topsoil (0-15cm) and ammonium (N-NH_4_^+^) in the respective soil layer in an experimental field at CIMMYT’s research station in Ciudad Obregon, North Mexico in the field season 2021/2022 (Exp. 3). Soil samples were taken from sowing rows at 7, 14 and 21 days after the second ammonium-N fertilizer application (DANF) and at harvest. ROELFS = parental line, control. CSMONO3B/3/CS/LE.RA//CS/4/CS ph/5/5*ROELFS = ROELFS line that carries the Lr#N-SA from *Leymus racemosus*. HIGH N: ammonium-N fertilizer application was split into 83 kg N ha^-1^ at sowing and 167 kg N ha^-1^ at booting stage. LOW N: ammonium-N fertilizer application as a single application of 83 kg N ha^-1^ at sowing. Error bars display the standard error of the mean of 3 replicated plots. Significant differences for the pairwise comparison for each sampling date separately is indicated for p-levels *<0.01, **<0.001 and ***<0.0001. n.s. = not significant.

In Exp 3 (Fig 3), nitrate in the row of ROELFS-Lr#N-SA compared to the ROELFS was 7% lower at 7 DANF (p= 0.0305), 37% lower at 14 DANF (p= 0.0026), and 24% lower (p=0.0002) at 21 DANF (HIGH N). The pattern was comparable at harvest with 34% lower topsoil NO_3_ ^-^ concentration in the BNI-wheat plots compared to the control (p=0.0024). Comparing HIGH N and LOW N of the respective lines always resulted in higher soil nitrate values in the plots that received N fertilizer twice, when comparing to the single dose application at sowing.

A significant NH_4_ ^+^ increase due to the 2^nd^ N fertilization in Exp 3 (Fig 3) was measured at 7 and 14 DANF but was absent at 21 DANF and at harvest. Significantly lower NH_4_ ^+^ concentrations were measured in the row of ROELFS-Lr#N-SA compared to ROELFS control two weeks after the second N split application (p=0.0067). However, this pattern could not be observed during the other three sampling dates (p=0.1546 for 7 DANF, p=0.1437 for 21 DANF and p=0.8090 at harvest).

Table 1 presents the nitrate amount in kg N–NO_3_ ^-^ ha^-1^ calculated from measured nitrate concentrations shown in Fig 1 and Fig 3. In Exp 1, for ROELFS-Lr#N-SA 8-9 kg N-NO_3_^-^ (+21%) was estimated in the row at 14 DANF, whereas in the furrow at 14 DANF, 16-17 kg N-NO ^-^ less was observed for ROELFS-Lr#N-SA. In Exp 3, nitrate amounts were 24 -36% lower in the sowing row. In Exp 3 nitrate amounts in plots of ROELFS-Lr#N-SA were 77 % and 73% lower at 33 days after the second N fertilization (Exp 1) measured in the sowing row and the furrow respectively.

**Table 1:** Topsoil (0-15cm) nitrate in kg N–NO_3_^-^ ha^-1^ calculated from measured nitrate concentrations shown in Fig. 1 and Fig. 3. Plots were planted with ROELFS = parental line, control, and CSMONO3B/3/CS/LE.RA//CS/4/CS ph/5/5*ROELFS = ROELFS line that carries the Lr#N-SA from *Leymus racemosus*. Exp 1 was conducted in 2019/20 and Exp 3 in 2021/2022. Plots were fertilized with 83 kg N ha^-1^ at sowing and 167 kg N ha^-1^ at booting stage (HIGH N), 60 days after sowing. Soil samples were collected from the sowing row and/or the furrow during different time points presented here in days after the second ammonium-fertilizer (DANF) split application.

| 2020 (Exp 1) | Sampling time | Nitrate-N in kg ha<sup>-1</sup> |  |  |
| --- | --- | --- | --- | --- |
|  |  | ROELFS Control | ROELFS+Lr#N-SA | Soil nitrate reduction or increase |
| Sowing row | 14 DANF | 41 | 50 | +9 |
|  | 33 DANF | 233 | 54 | -179 |
| Furrow | 14 DANF | 36 | 53 | -17 |
|  | 33 DANF | 195 | 53 | -142 |
| 2022 (Exp 3) | Sampling time | Nitrate-N in kg ha<sup>-1</sup> |  |  |
|  |  | ROELFS Control | ROELFS+Lr#N-SA | Soil nitrate reduction or increase |
| Sowing row | 14 DANF | 72 | 46 | -16 to -17 |
|  | 21 DANF | 76 | 59 | -18 to -19 |
|  | harvest | 37 | 25 | -12 to -13 |

### Leaf nitrate

Two months after sowing and after the application of 83 kg of ammonium-N in Exp 1, the plants reached the booting stage. Flag leaves nitrate concentration at booting (Fig 4, before second N application) was significantly higher in ROELFS-Lr#N-SA compared to the control (p<0.0001). Continuous monitoring of the nitrate concentration in flag leaves after an additional 167 kg ammonium-N ha^-1^ application (HIGH N) and flood-irrigation (HIGH N and LOW N) showed higher values in ROELFS-Lr#N-SA compared to ROELFS (Fig 5) at 2, 5, 8 and 12 days after the second N (DANF) split application. The significantly increased flag leaf nitrate concentration of the translocation line compared to the control was also an observation in 3 out of 4 sampling dates in the LOW N plots.

**Fig 4:**
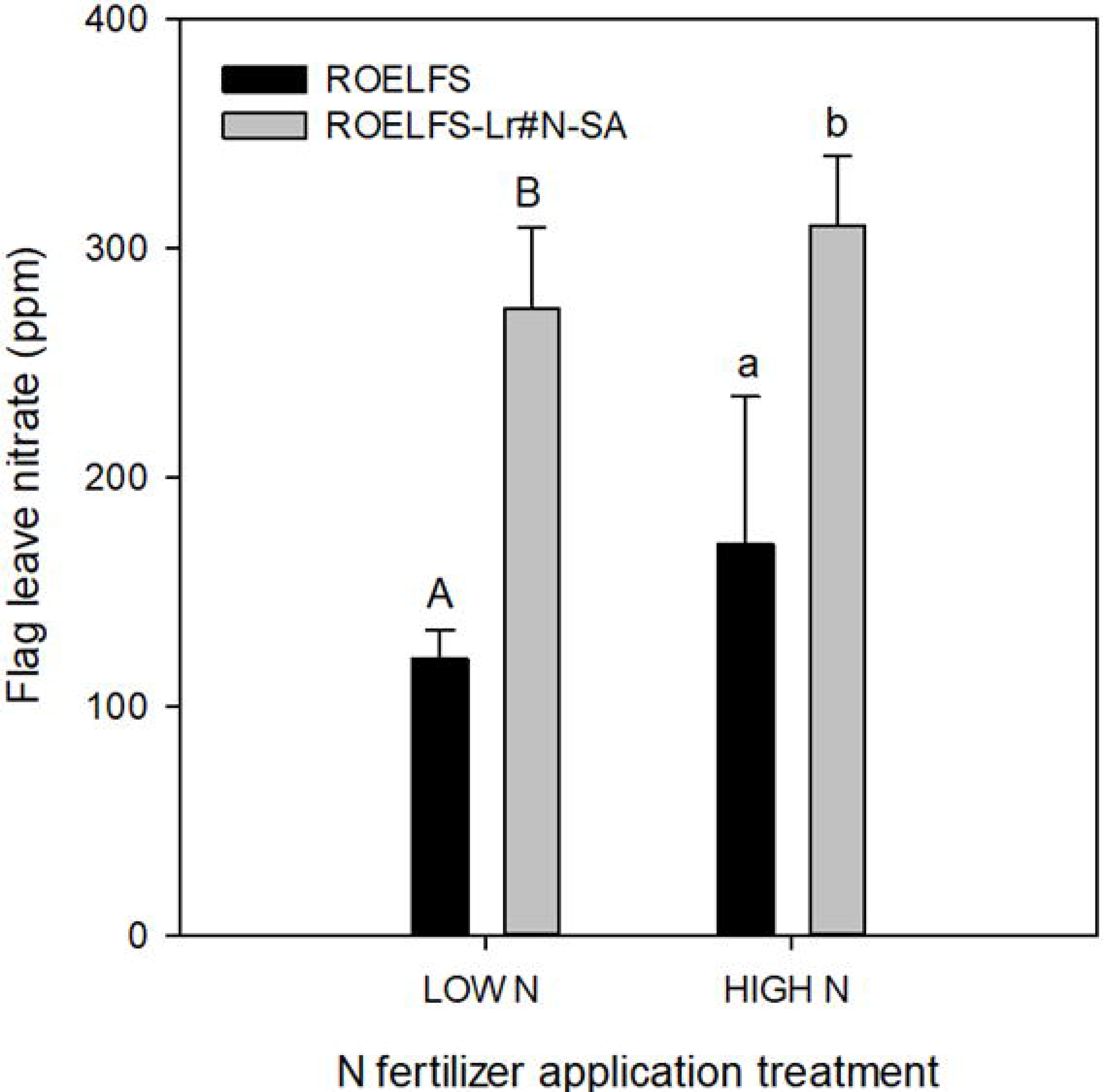
Nitrate concentration measured in flag leaves at booting of ROELFS parental and ROELFS-Lr#N-SA (BC_5_) lines at CIMMYT’s research station in Ciudad Obregon 2019/2020 (Exp. 1) at 60 days after sowing, just before the second N fertilizer application for HIGH N plots. LOW N = single application at sowing 83 kg N ha^-1^ in form of ammonium sulphate. HIGH N = split N application with ammonium sulphate 83 kg N ha^-1^ at sowing and 167 kg N ha^-1^ at booting. The pairwise comparison was conducted for HIGH N (capital letters) and LOW N (lowercase letters) separately. Significant differences between the two lines (p<0.0001) are indicated with different letters.

**Fig 5:**
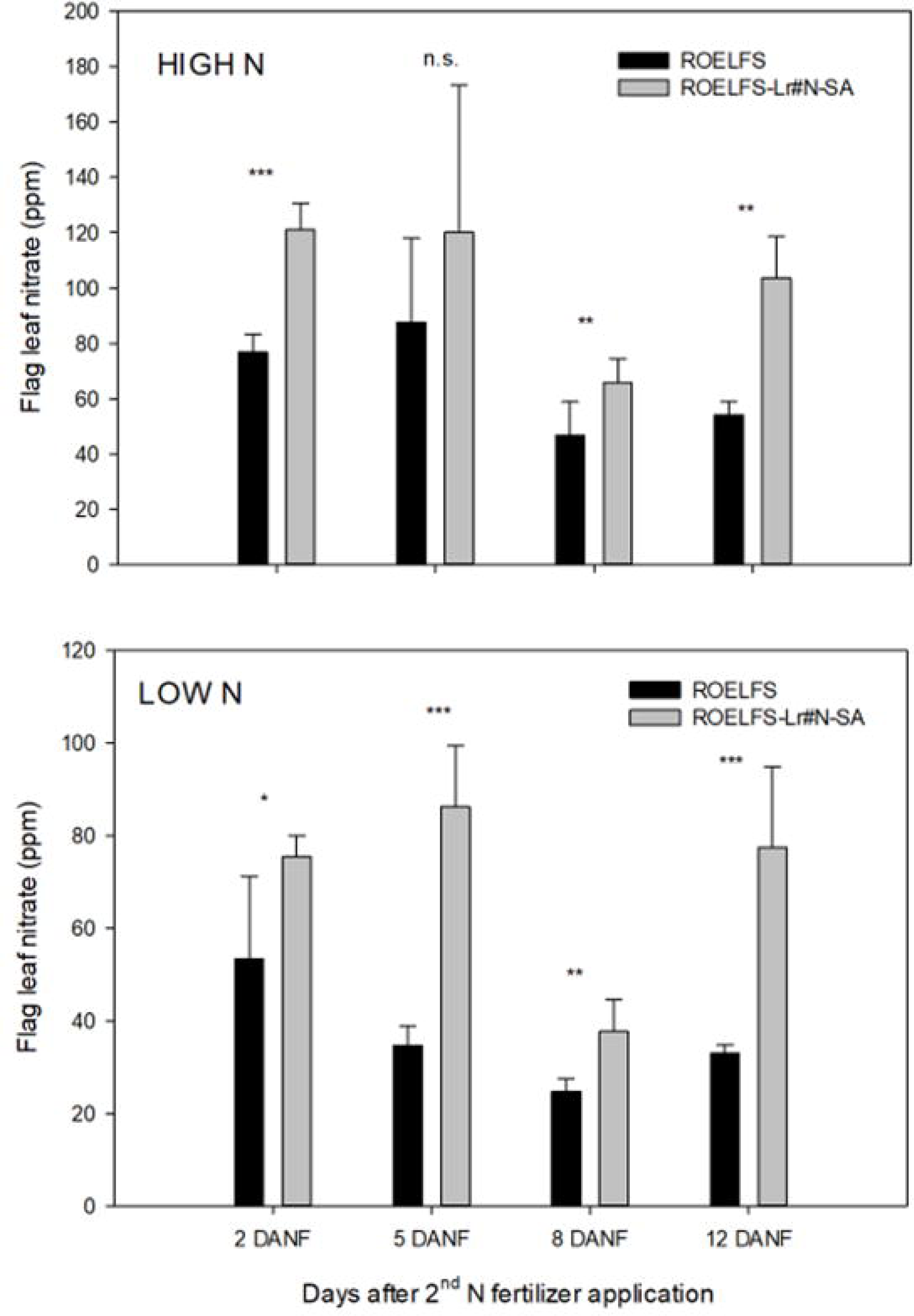
Nitrate concentration (in ppm NO_3_) measured in flag leaves at booting of ROELFS parental and ROELFS-Lr#N-SA (BC_5_) lines in Obregon 2019/2020 (Field trial 1) in HIGH N and LOW N (no second N fertilizer application) at 2, 5, 8 and 12 days after the second N fertilizer (DANF) application. Significant differences for the pairwise comparison for each sampling date separately are indicated for p-levels *<0.01, **<0.001 and ***<0.0001. n.s. = not significant.

### Potential nitrification rates

The results from soil incubation (Exp 2 and Exp 3) expressed as potential nitrification rates (PNRs) are shown in Fig 6. Soil taken periodically from the HIGH N plots after the second N fertilization always showed higher in-vitro nitrification rates compared to the plot that did not receive the second ammonium dose in the field (p<0.0001). Significantly less nitrate was produced over time in soil taken one week after the second N application for the translocation line. This observation was consistent with a PNR of 2.04 (in mg N-NO_3_ ^-^ per kg dry soil day^-1^) for the ROELFS-Lr#N-SA line versus a PNR of 2.81 for the control in Exp 2 (p=0.0485) and PNRs of 1.86 for ROELFS-Lr#N-SA compared to a 2.74 for ROELFS control in Exp 3 (p=0.0492). However, PNRs of soil that was sampled later (14 and 21 DANF) did not differ significantly between ROELFS-Lr#N-SA and control lines in both experiments (p>0.05). Furthermore, no significant different PNRs were measured in incubated soil from the ROELFS pair form LOW N plots.

**Fig 6:**
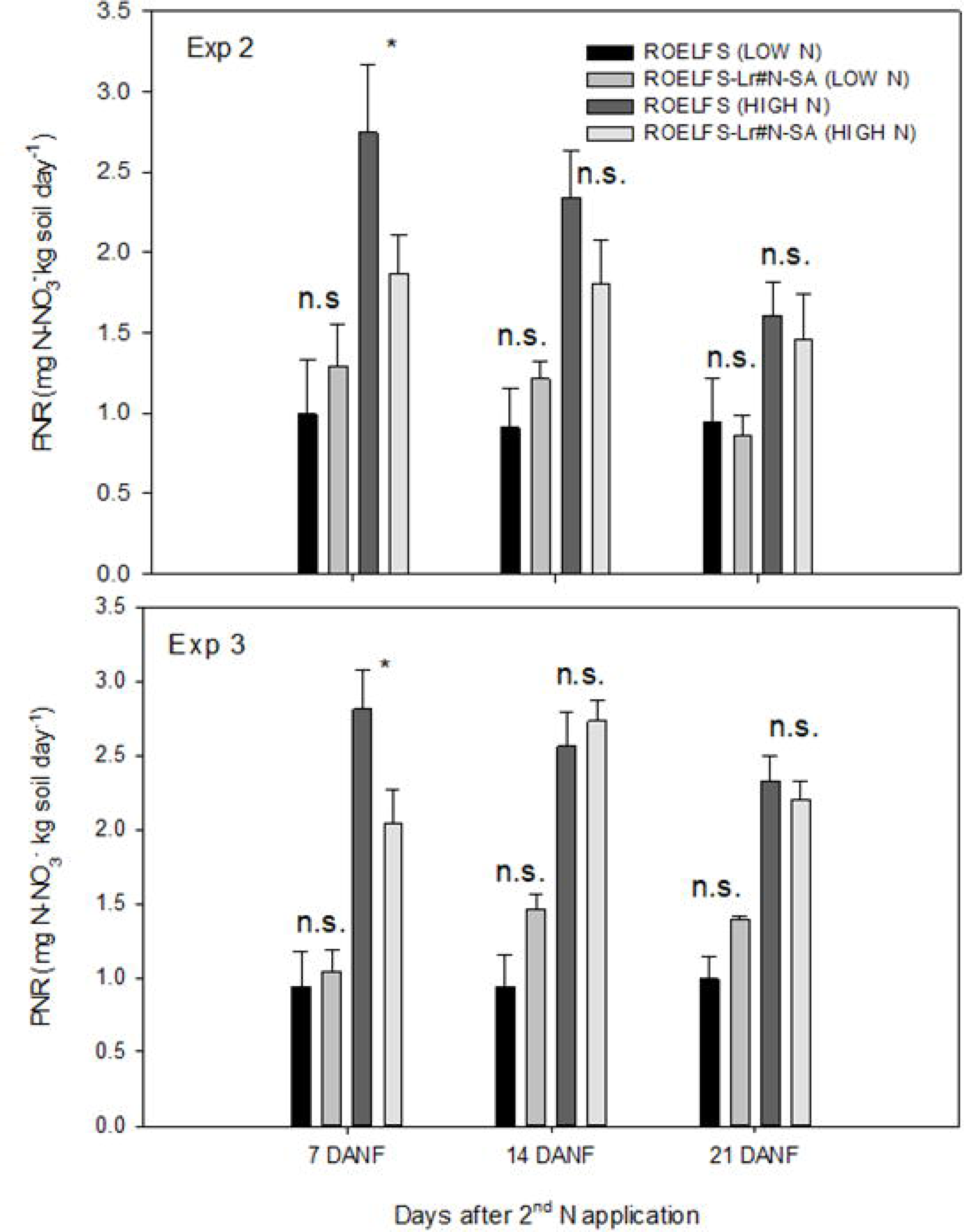
Potential nitrification rates (PNRs) measured with a soil-slurry method (mg N-NO_3_ per kg dry soil day). Soil samples were collected from the topsoil (0-15cm) of the sowing rows at 7, 14 and 21 days after the second N fertilizer application (DANF) in 2020/2021 in Exp. 2 and 2021/2022 in Exp.3. Significant differences for the pairwise comparison for each sampling date separately are indicated for p-levels *<0.05. n.s. = not significant.

### Grain yield and yield parameters Experiment 1

Grain yield (Table 2) was significantly lower for the ROELFS-Lr#N-SA line (p=0.0004) in Exp 1 and a significant grain yield increase was not observed due to additional N application (p=0.1051). The interaction of the genotype*N rate had no effect on grain yield (p=0.4182). Biomass for the translocation line was also significantly lower compared to the control line (p=0.0121). Again, an additional N application did not increase biomass (p=0.1789). The harvest index was also lower for the translocation line (p=0.0069), regardless of additional N fertilizer (p=0.8953). The number of spikes per square meter did not differ among the lines (p=0.4944) nor among the N rate treatments (p=0.4401) or the interaction of both (p=0.7560). However, less grains per spikes were observed for the BNI line (p=0.0290) and the LOW N plots (p=0.0438). Grains per spike were also lower for ROELFS-Lr#N-SA (p=0.0009) compared to ROELFS but not affected by the N fertilizer (p=0.6536). The translocation line reached anthesis significantly later than the control line (p=0.0036), which was affected by the N fertilizer treatment (p=0.4458). Maturity was mainly influenced by genotype (p=0.0046) but also by the N fertilizer (p=0.0240). ROELFS-Lr#N-SA was later maturing than ROELFS control and N application increased maturity of both lines. The thousand kernel weight did not differ significantly among the lines (p=0.2297) or the N rate (p=0.9585).

**Table 2:**
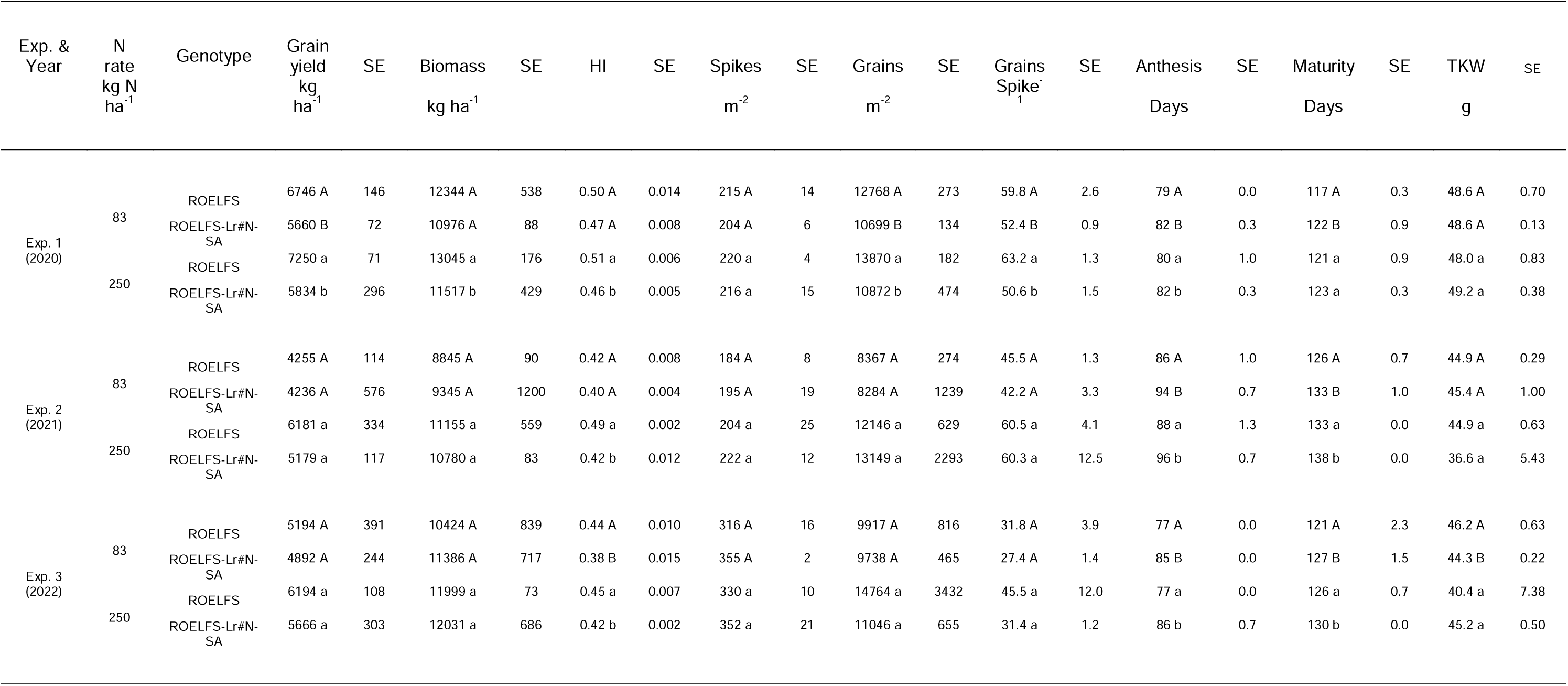
Grain yield, biomass and yield components (HI=harvest index; TKW=thousand kernel weight). N rate 250 = 83 kg N ha^-1^ at sowing and 167 kg N ha^-1^ at booting (HIGH N). N rate 83 = 83 kg N ha^-1^ at sowing, no N fertilizer at booting (LOW N). Pairwise-comparison for ROELFS-Lr#N-SA and ROELFS was conducted separately for each N rate treatment (250 or 83) and year (2020, 2021 or 2022). Exp. 1 was conducted in 2020 in one field whereas Exp.2 and Exp.3 were based on two consecutive years (2021 and 2022) in another field. The same letters indicate no significant difference between the mean values of the two lines. SE=standard error of the mean from three field replications. *Data for Exp 2 under 250 N were published already as reference data by Bozal-Leorri et al. 2022*.

| Exp. & Year | N rate kg N ha<sup>-1</sup> | Genotype | Grain yield kg ha<sup>-1</sup> | SE | Biomass kg ha<sup>-1</sup> | SE | HI | SE | Spikes m<sup>-2</sup> | SE | Grains m<sup>-2</sup> | SE | Grains Spike<sup>-1</sup> | SE | Anthesis Days | SE | Maturity Days | SE | TKW g | SE |
| --- | --- | --- | --- | --- | --- | --- | --- | --- | --- | --- | --- | --- | --- | --- | --- | --- | --- | --- | --- | --- |
| Exp. 1 (2020) | 83 | ROELFS | 6746 A | 146 | 12344 A | 538 | 0.50 A | 0.014 | 215 A | 14 | 12768 A | 273 | 59.8 A | 2.6 | 79 A | 0.0 | 117 A | 0.3 | 48.6 A | 0.70 |
|  |  | ROELFS-Lr#N-SA | 5660 B | 72 | 10976 A | 88 | 0.47 A | 0.008 | 204 A | 6 | 10699 B | 134 | 52.4 B | 0.9 | 82 B | 0.3 | 122 B | 0.9 | 48.6 A | 0.13 |
|  | 250 | ROELFS | 7250 a | 71 | 13045 a | 176 | 0.51 a | 0.006 | 220 a | 4 | 13870 a | 182 | 63.2 a | 1.3 | 80 a | 1.0 | 121 a | 0.9 | 48.0 a | 0.83 |
|  |  | ROELFS-Lr#N-SA | 5834 b | 296 | 11517 b | 429 | 0.46 b | 0.005 | 216 a | 15 | 10872 b | 474 | 50.6 b | 1.5 | 82 b | 0.3 | 123 a | 0.3 | 49.2 a | 0.38 |
| Exp. 2 (2021) | 83 | ROELFS | 4255 A | 114 | 8845 A | 90 | 0.42 A | 0.008 | 184 A | 8 | 8367 A | 274 | 45.5 A | 1.3 | 86 A | 1.0 | 126 A | 0.7 | 44.9 A | 0.29 |
|  |  | ROELFS-Lr#N-SA | 4236 A | 576 | 9345 A | 1200 | 0.40 A | 0.004 | 195 A | 19 | 8284 A | 1239 | 42.2 A | 3.3 | 94 B | 0.7 | 133 B | 1.0 | 45.4 A | 1.00 |
|  | 250 | ROELFS | 6181 a | 334 | 11155 a | 559 | 0.49 a | 0.002 | 204 a | 25 | 12146 a | 629 | 60.5 a | 4.1 | 88 a | 1.3 | 133 a | 0.0 | 44.9 a | 0.63 |
|  |  | ROELFS-Lr#N-SA | 5179 a | 117 | 10780 a | 83 | 0.42 b | 0.012 | 222 a | 12 | 13149 a | 2293 | 60.3 a | 12.5 | 96 b | 0.7 | 138 b | 0.0 | 36.6 a | 5.43 |
| Exp. 3 (2022) | 83 | ROELFS | 5194 A | 391 | 10424 A | 839 | 0.44 A | 0.010 | 316 A | 16 | 9917 A | 816 | 31.8 A | 3.9 | 77 A | 0.0 | 121 A | 2.3 | 46.2 A | 0.63 |
|  |  | ROELFS-Lr#N-SA | 4892 A | 244 | 11386 A | 717 | 0.38 B | 0.015 | 355 A | 2 | 9738 A | 465 | 27.4 A | 1.4 | 85 B | 0.0 | 127 B | 1.5 | 44.3 B | 0.22 |
|  | 250 | ROELFS | 6194 a | 108 | 11999 a | 73 | 0.45 a | 0.007 | 330 a | 10 | 14764 a | 3432 | 45.5 a | 12.0 | 77 a | 0.0 | 126 a | 0.7 | 40.4 a | 7.38 |
|  |  | ROELFS-Lr#N-SA | 5666 a | 303 | 12031 a | 686 | 0.42 b | 0.002 | 352 a | 21 | 11046 a | 655 | 31.4 a | 1.2 | 86 b | 0.7 | 130 b | 0.0 | 45.2 a | 0.50 |

### Experiment 2

After the first year (2021) at the second field site, the grain yield was significantly higher for wheat in HIGH N compared to LOW N plots (p=0.0132), but no difference was observed between the lines in each plot (p=0.1865) (Table 2). The interaction of N rate and the wheat line was not significant (p=0.6906). No differences for the biomass were indicated according (p=0.1266) to the N rate treatment, p=0.4669 for the line comparison and the respective interaction p=0.4952. The harvest index was significantly lower at LOW N compared to HIGH N plots (p=0.0282) and lower for the translocation line compared to the control (p=0.0008). Both anthesis and days to maturity of the translocation line were significant later than the control in both N treatments (p<0.0001). Plants in the HIGH N plots also matured later (p=0.0185) than the ones that received the LOW N doses. Grain N% was higher in high N rate (p<0.0001 for both NIR and DUMAS method) but no difference was observed between genotypes within each N rate. (p=0.6444 (NIR) and p=0.3561 (DUMAS). N uptake was significantly increased by the second N split application (p<0.0001) but no difference was observed between the two wheat lines (p=0.3456).

### Experiment 3

Grain yield depended on the N rate (p=0.0132) but not on the genotype (p=0.1865) or the interaction (p=0.6906). There was no significant difference in the biomass between the N rate treatments (p=0.2583), the two lines (p=0.3553) or the interaction of both (p=0.4952) (Table 2). The harvest index differed significantly between the N treatments (p=0.0282) and the lines (p=0.0008). The N rate treatment and the genotype had no effect on the number of counted spikes per area (p=0.7155). The ROELFS-Lr#N-SA line reached anthesis significantly later compared to ROELFS-control (p<0.0001), an observation less dependent from the N rate (p=0.0805). There was a significant difference for days to maturity at the different N levels (p=0.0185) as well as between the lines (p=0.0065) but no interaction of both factors on maturity was noted (p=0.3198).

### Grain quality (Table 3, Exp 2 and Exp 3)

Grain N concentration measured with NIR was affected significantly by the N fertilizer application rate (p<0.0001) but there was no difference between the lines (p=0.6444). N% in grains measured with the DUMAS methodology confirmed the NIR results for N level (p<0.0001) and no significant difference for the line comparison (p=0.3561). N uptake was higher than under 250 N compared to 83 N (p<0.0001) but not different among the lines (p=0.3456). Grain protein did not differ significantly among the lines (p=0.8950) but was higher for the HIGH N treatment (p=0.0576).

The comparison of the wheat lines separately for each year and N treatment did not result in any significant difference (Table 3).

**Table 3:** Grain quality (grain protein, grain N concentration d) measured with near infrared analysis (NIR) and a dry combustion method. Grain N uptake was calculated by multiplying grain yield ha^-1^ with the grain N concentration (in %). N rate 250 = 83 kg N ha^-1^ at sowing and 167 kg N ha^-1^ at booting (HIGH N). N rate 83 = 83 kg N ha^-1^ at sowing, no N fertilizer at booting (LOW N). Pairwise-comparison for ROELFS-Lr#N-SA and ROELFS was conducted separately for each N rate treatment (250 or 83) and year (2020, 2021 or 2022). The same letters indicate no significant difference between the mean values of the two lines. SE=standard error of the mean from three field replications.

| Year | N rate<br>kg N ha<sup>-1</sup> | Genotype | Grain Protein<br>% | SE | Grain N% | SE | Grain N uptake<br>kg N ha<sup>-1</sup> | SE |
| --- | --- | --- | --- | --- | --- | --- | --- | --- |
| Exp. 2 (2021) | 83 | ROELFS | 8.1 A | 0.31 | 1.39 A | 0.053 | 56.9 A | 1.0 |
|  |  | ROELFS-Lr#N-SA | 8.7 A | 0.33 | 1.48 A | 0.057 | 61.1 A | 9.8 |
|  | 250 | ROELFS | 11.8 a | 0.44 | 2.02 a | 0.075 | 120.9 a | 11.1 |
|  |  | ROELFS-Lr#N-SA | 11.6 a | 0.23 | 1.98 a | 0.040 | 99.4 a | 3.9 |
| Exp. 3 (2022) | 83 | ROELFS | 8.8 A | 0.48 | 1.52 A | 0.083 | 77.7 A | 9.2 |
|  |  | ROELFS-Lr#N-SA | 9.1 A | 0.30 | 1.56 A | 0.052 | 75.2 A | 6.2 |
|  | 250 | ROELFS | 11.9 a | 0.17 | 2.04 a | 0.029 | 123.9 a | 3.9 |
|  |  | ROELFS-Lr#N-SA | 12.4 a | 0.52 | 2.13 a | 0.090 | 118.0 a | 4.7 |

In both years, the genotype had a significant effect on test weight, with the ROELFS-Lr#N-SA lines generally showing a 1.4 to 2.6% decrease in test weight values compared to the control. In general, no highly significant effect from either the N treatment or the genotype was observed for thousand kernel weight. However, both factors affected the grain hardness values in both years, with the treatment generally having a stronger influence on this trait. In fact, lines grown under 250 kg N ha ¹ showed a general 9 to 17% increase in grain hardness compared to lines grown under 83 kg N ha ¹. Slight but significantly lower hardness values were also observed for the control compared to the ROELFS-Lr#N-SA line. Flour yield was not influenced by the genotype but was affected by the treatment, with lines grown under the higher N fertilization conditions generally showing higher flour yield values (year 2022 average = 70.87%; year 2021 average = 71.51%) compared to those grown under lower N fertilization (year 2022 average = 68.20%; year 2021 average = 69.49%) (Table 4).

**Table 4:** Average grain, milling, gluten, dough and ened-use quality parameters evaluated in the BNI (ROELFS-Lr#N-SA) line and its relative control variety (ROELFS) across two different years (2022 and 2021) at two different N levels (250 and 83 kg N ha^-1^). Different letters indicate a significant difference between the mean values recorded within one year. Avg = Average; SE = Standard Error; TW=Test Weight; TKW = Thousand Kernel Weight; GRNHRD = Grain Hardness; FLRYLD = Flour Yield; FLRSDS = SDS Sedimentation Volume; ALVW = Alveograph W; ALVPL = Alveograph P/L; LOFVOL = Loaf Volume; CRMSTR = Crumb Structure.

| Year | N Rate kg<br>N ha<sup>-1</sup> | Genotype | Grain and Milling Quality Parameters |  |  |  |  |  |  |  |
| --- | --- | --- | --- | --- | --- | --- | --- | --- | --- | --- |
|  |  |  | TW (kg/hL) |  | TKW (g) |  | GRNHRD (HI) |  | FLRYLD (%) |  |
|  |  |  | Avg | SE | Avg | SE | Avg | SE | Avg | SE |
| 2022 | 250 | ROELFS-Lr#N-SA | 78.82 b | 0.14 | 48.15 | 0.34 | 65.72 a | 0.46 | 70.58 a | 0.36 |
|  |  | ROELFS | 80.98 a | 0.1 | 49.23 | 0.29 | 63.42 ab | 0.19 | 71.15 a | 0.18 |
|  | 83 | ROELFS-Lr#N-SA | 78.64 b | 0.06 | 47.82 | 0.24 | 61.07 b | 0.85 | 68.62 b | 0.21 |
|  |  | ROELFS | 80.53 a | 0.21 | 48.71 | 0.07 | 57.17 c | 0.28 | 67.77 b | 0.27 |
| 2021 | 250 | ROELFS-Lr#N-SA | 78.33 c | 0.09 | 46.72 b | 0.32 | 68.05 a | 0.27 | 70.09 b | 0.22 |
|  |  | ROELFS | 79.48 ab | 0.1 | 48.65 ab | 0.24 | 61.39 ab | 0.25 | 72.92 a | 0.20 |
|  | 83 | ROELFS-Lr#N-SA | 78.68 bc | 0.17 | 49.34 a | 0.42 | 58.34 bc | 1.71 | 69.67 b | 0.36 |
|  |  | ROELFS | 79.65 a | 0.15 | 49.73 a | 0.35 | 52.41 c | 1.27 | 69.31 b | 0.32 |

| Year | N Rate kg<br>N ha<sup>-1</sup> | Genotype | Gluten and Dough Rheological Properties |  |  |  |  |  | Bread Making Quality |  |  |  |
| --- | --- | --- | --- | --- | --- | --- | --- | --- | --- | --- | --- | --- |
|  |  |  | FLRSDS (mL) |  | ALVW (J10<sup>-4</sup>) |  | ALVPL |  | LOFVOL (cm<sup>3</sup>) |  | CRMSTR |  |
|  |  |  | Avg | SE | Avg | SE | Avg | SE | Avg | SE | Avg | SE |
| 2022 | 250 | ROELFS-Lr#N-SA | 18.00 a | 0.19 | 284.00 b | 7.02 | 1.63 b | 0.17 | 741.67 a | 2.2 | 4.00 a | 0.00 |
|  |  | ROELFS | 17.58 a | 0.11 | 298.67 ab | 6.01 | 1.34 b | 0.15 | 761.67 a | 2.2 | 4.00 a | 0.00 |
|  | 83 | ROELFS-Lr#N-SA | 12.92 b | 0.3 | 256.67 b | 4.48 | 3.14 a | 0.17 | 645.00 b | 12.6 | 3.00 b | 0.00 |
|  |  | ROELFS | 14.50 b | 0.38 | 346.67 a | 13.47 | 3.17 a | 0.24 | 680.00 b | 5.0 | 3.33 b | 0.17 |
| 2021 | 250 | ROELFS-Lr#N-SA | 14.50 a | 0.12 | 255.67 a | 3.53 | 1.75 b | 0.21 | 691.67 ab | 6.0 | 4.00 a | 0.00 |
|  |  | ROELFS | 15.00 a | 0.07 | 226.67 ab | 9.11 | 1.19 b | 0.04 | 713.33 a | 6.8 | 4.00 a | 0.00 |
|  | 83 | ROELFS-Lr#N-SA | 9.42 b | 0.4 | 192.67 b | 7.37 | 4.66 a | 0.5 | 536.67 c | 27.2 | 2.33 b | 0.17 |
|  |  | ROELFS | 10.58 b | 0.04 | 261.33 a | 9.39 | 3.82 a | 0.15 | 591.67 bc | 14.2 | 3.00 b | 0.00 |

Results from the SDS sedimentation volume test showed that, independently of the genotype, lines grown under higher N concentrations were associated with 30 to 47% higher SDS sedimentation volumes, indicative of generally better overall gluten quality. Gluten strength, as expressed by the alveograph W value, was generally more influenced by the genotype than by the treatment, with ROELFS exhibiting slightly stronger gluten (year 2022 average = 322.67 J10^-4^; year 2021 average = 244 J10^-4^) compared to the ROELFS-Lr#N-SA line (year 2022 average = 270.34 J10^-4^; year 2021 average = 224.17 J10^-4^). In contrast, dough balance (dough tenacity/dough extensibility as expressed by the alveograph P/L value) appeared to be influenced only by the treatment, with lines grown under low N regimes being associated with a more tenacious and less balanced dough (year 2022 average = 3.16; year 2021 average = 4.24) compared to lines grown under 250 kg N ha ¹ (year 2022 average = 1.49; year 2021 average = 1.47). A similar trend was observed for the bread-making quality of the evaluated lines, where the N provided to the plant during the growing cycle was the main factor influencing the results. Indeed, 13 to 25% higher loaf volumes as well as better crumb structure were observed in the lines grown under higher N fertilization regimes compared to lines grown with lower N inputs (Table 4).

To better understand the reasons behind variation in gluten strength, the ROELFS-Lr#N-SA and ROELFS lines were evaluated for their glutenin profile using SDS-PAGE. This analysis indicated that, in general, both lines shared the same glutenin profile (*Glu-A1b*; *Glu-B1i*; *Glu-D1d*; *Glu-A3b*; *Glu-B3h*; and *Glu-D3b*). However, it was noted that all the ROELFS-Lr#N-SA lines also carried a second allele at the *Glu-B1* locus (*Glu-B1a*) and at the *Glu-A3* locus (*Glu-A3c*), and that they had a different allele at the *Glu-D3* locus compared to ROELFS (*Glu-D3c* instead of *Glu-D3b*).

Overall, based on the observed wheat quality data, the nitrogen treatment appeared to be the main factor influencing these values, even though critical characteristics such as test weight, grain hardness and gluten strength appeared to be negatively influenced by the BNI translocation.

## Discussion

### BNI as a seed-based solution to reduce N environmental pollution in high production wheat systems

In the Cajeme district of the Yaqui Valley, irrigated spring wheat is produced on an area of 150,000–180,000 ha (Fischer et al. 2014); 19% of the average applied 250 kg N per ha was, and likely still is leached directly from the field during every season (Grahmann et al. 2018). This would result in per season 7125 to 8550 t of leached nitrate-N into groundwater bodies, drains and rivers flowing into the Gulf of California. The approximate economic loss (if the price for one 1 t of urea is assumed 1000 U.S. dollars) of this N Loss sums up to more than 8.5 million USD annually. One would need to add the indirect costs of the pollutive effects to the assessment.

An ex-ante impact study (Leon et al. 2022) predicted that BNI-wheat with an assumed 40% nitrification inhibition (developed by 2050) could lead to a 15% reduction, in N fertilization, 16% reduction in LC-GHG emissions and 17% improvement in NUE at farm level; a reduction in GHG emissions (e.g., N_2_O) between 7.3 and 9.5% across the suitable wheat-harvested area worldwide could be achieved with 30% and 40% nitrification inhibition, respectively. We observed a reduction of nitrate in the topsoil within a range of 32-77% with spatial and temporal variation, hypothesis 1) could therefore be confirmed. Whilst we did not investigate the legacy or carry over effect of nitrification between cropping seasons, it has been shown that a residual BNI effect exists, although persistence declines with time (Karwat et al. 2017). Our data suggests that BNI lines could deliver a potential reduction of nitrification (27-32% lower PNRs, Fig 6) estimated by Leon et al. (2022). Furthermore, the BNI effect was observed in actual harvest (Fig 3). These encouraging results call for further multi-location testing to estimate the scale of this net positive effect on the environment in wheat growing areas globally (Braun et al. 2010). A linear reduction of the N fertilizer amounts in irrigated spring wheat system as in the Yaqui Valley could reduce N_2_O emissions exponentially (Millar et al., 2018).

### BNI expression in alkaline soils

Field observations of BNI model plants like *Brachiaria humidicola* (Subbarao et al. 2009) confirmed effective inhibition of soil nitrification in neutral pH conditions (pH 7.2). For *sorghum bicolor,* rhizosphere pH influenced release of hydrophilic-BNIs, but not of hydrophobic-BNIs (Di et al. 2018). The BNI activity of ROELFS-Lr#N-SA was equivalent to 162 allylthiourea units (ATU), versus 86 ATU for ROELFS-Control (Subbarao et al. 2021). Whilst the metabolic fingerprinting of root exudates from BNI-wheat lines are pending, preliminary results suggest that the main BNIs are hydrophilic (pers. comm Subbarao). That ROELFS-Lr#N-SA releases hydrophilic BNIs is also implied by the BNI effect observed in the furrow (40cm from sowing rows) of this study (Table 1). Egenolf et al. (2021) conducted root exudate collection studies with *Brachiaria humidicola* and confirmed higher release of BNIs under low pH in the trap solution, though cations, e.g. potassium (K^+^), remain overlooked as a potential trigger for BNI release. It can be assumed that the active release of hydrophilic BNIs is increased by low pH (soils or hydroponic media). Furthermore, the release of hydrophobic BNI compounds in high pH soils needs to be investigated.

Subbarao et al. (2021) had reported on BNI activity in Japanese acidic soil (pH 5.6) where BNI was triggered by three N split applications of ammonium sulphate solutions. Our research made the case for a protocol in alkaline soils (pH 8.6-8.7), based on conventional ammonium sulphate fertilizer applied as one third of the dose at sowing and two thirds at booting stage. The BNI effect could be confirmed in the form of significant lower PNRs in two consecutive years in soil samples taken 7 days after the second N application (Fig 6). But the PNRs in soil taken 14 and 21 days after NH_4_ ^+^ field addition did not differ between the BNI and the control line. This confirms partly our hypothesis 2). These observations suggest a strong release of BNIs triggered shortly after ammonium addition (Subbarao et al. 2007), despite high soil pH.

Although the significant persistence of the BNI effect could not be confirmed with the in vitro soil slurry method, the in-situ nitrate monitoring confirmed strong BNI expression for weeks after N fertilization (Fig 1 and Fig 3). The observation that soil nitrate was reduced more than 30% even after harvest (Fig 3 and Table 1) indicates that apart from active BNI release, a considerable residual effect contributed to BNI expression. We propose that PNRs from *in vitro* observations are useful indicators in addition to in situ N measurements for BNI expression, though the discrepancy between the field soil and soil slurry might bias for or against certain nitrifying groups (Hazard et al. 2021).

### Higher plant nitrate uptake as secondary beneficial effect due to the Lr#N translocation to reduce soil nitrate

Low leaf nitrate has been used as an indicator for high BNI expression of the tropical pasture *Brachiaria humidicola* (Karwat et al. 2019) and adapted to study N dynamics of BNI-wheat in pot studies (Bozal-Leorri et al. 2022). The higher nitrate values found for the spring wheat translocation line (Fig 4 and 5) are in contrast with other studies’ findings. Lower soil nitrate of ROELFS-Lr#N-SA was a consistent observation under high N input management (Fig 1 and Fig 3) within the presented field studies but was not reflected by lower leaf nitrate concentrations of the respective wheat lines. Hypothesis 4), that lower soil nitrate is reflected in lower plant tissue nitrate could therefore not be confirmed. Since this was a frequent observation under both high and low N, higher plant nitrate uptake could be an additional trait transferred to ROELFS by the Lr#N-SA. Fast and efficient uptake may play an important role in the measured soil nitrate reduction in the root zone soil. Furthermore, higher N loading of the sink at booting could potentially have a positive effect on grain formation (Fradgley et al. 2021), although our observation in 2020 (Table 2) could not confirm that. Subbarao et al. (2021) mentioned increased mineralization by translocation lines, something not expected due to BNI per se. We suggest to further collect further observations if the Lr#N-SA translocation in elite lines increases nitrate uptake and assimilation and if this constitutes a complementary trait to BNI in terms of soil nitrate reduction and consequently its loss.

### Soil ammonium patterns under BNI

Soil ammonium concentrations of the translocation line at the root zone and in the furrow of the respective plots were around 20 ppm lower compared to the NH_4_ ^+^ concentrations (Fig 2) compared to the control line in (Exp 1), This observation did not match with our hypothesis 3) (e.g. significantly higher NH_4_ ^+^ due to a decelerated nitrification). It remains unclear what is behind those lower concentrations. Ghosh et al. (2024) surmised BNI release activity due to the observed high ammonium retention of *Sorghum halepense*.

Lower ammonium in soil of the translocation line measured around four weeks (Fig 2 A – HIGH N) after the application of ammonium-N in Exp 1 was observed. In Exp 2 the lower ammonium observation in the topsoil of the BNI-line was confirmed 14 days after N dressing. In all other comparisons, ammonium soil N did not differ between the wheat lines. We have no indication that the N was lost in any other way, but BNI has been shown to increase microbial immobilization (Egenolf et al. 2022). Issifu et al. (2024) tested a range of BNIs and observed both synergistic and antagonistic effects on soil N dynamics. We concluded that release of BNIs may have stimulating on heterotrophic microbes that take up available mineral N. However, N that is immobilized in microbial biomass cannot be nitrified or denitrified. A secondary effect of BNIs on other microbial groups and soil mineral N availability and potential negative tradeoffs for the plants’ N nutrition (Kuppe and Postma, 2024) need to be investigated.

### Ammonium N fertilizer as a trigger for BNI expression

BNI expression in terms of reduced soil nitrate in the root zone and between sowing beds depended on the additional application of ammonium fertilizer (Fig 1, 2, 3 and Table 1). This observation was reported for “BNI-wheat” (Subbarao et al, 2021, Bozal-Leorri et al. 2022) but also for other BNI grasses (Karwat et al. 2018, 2019) or crops with confirmed BNI activity like Sorghum (Zhang et al. 2022). The expression of the effect was significant in microcosm slurry studies in the laboratory (Fig 6) taken one week after the N application but absent in soil from plots that did not receive a second N dose. That ammonium application is needed to express BNI was also indicated by frequent NO_3_ ^-^ monitoring in-situ/field (Fig 1) that showed a significant expression in the field two and three weeks after the second ammonium sulphate split application (Fig 3) with a constant pattern also observed at harvest. We confirm hypothesis 5, e.g. that ammonium addition to the system is necessary to trigger both the BNI expression from the plant and a reaction from the soil nitrifier community.

### Further improvement of elite wheat lines with BNI expression by breeding

The Lr#N-SA showed benefits on the environmental rather than on the agronomic side when introduced into ROELFS. In contrast, other field observations at CENEB (Year 2020) reported 13.8% of grain protein for ROELFS-Lr#N-SA compared to 13.2% for the control (Subbarao et al. 2021, Table S8a). The delayed maturity of the BNI-ROELFS by 3 days (Subbarao et al. 2021, Table S9) is in line with our current observation of 4-5 days delay in maturity (Table 2). Delaying anthesis and related senescence can positively impact wheat grain yields particularly under low N (Gaju et al. 2011). This needs to be investigated across a greater number of Lr#N elite wheat lines to come to a better understanding.

Gene introduction from wild wheat relatives is often accompanied by compromises in yield potential (Emebiri et al. 2021). Although our observation confirms yield stability in two of three years, but the lower HI for ROELFS-Lr#N-SA was a consistent observation. Therefore, hypothesis 6 (no negative effects of Lr#N-SA on yield potential quality traits) cannot be fully confirmed. It is critical to understand why a significant nitrification inhibition effect and an increased nitrate uptake did not translate into improved HI and total N amount in grains at harvest. We also suggest conducting more backcrosses of the Lr#N-SA line with ROELFS to improve its yield stability and yield quality.

Additionally, by comparing the quality profiles of the two lines, the general trend was that the line with the translocation exhibited lower test weight and gluten strength, and slightly lower grain hardness values compared to the control. While the reduction in test weight may be linked to the lower yield observed in some cases (Table 2), the lower gluten strength (Table 4) warrants further research.

Based on the glutenin profile analysis, the ROELFS-Lr#N-SA and ROELFS lines appear to have almost identical profiles, which is expected given the high number of backcrosses and the fact that the BNI translocation is not located on any of the major chromosomes (group 1 and group 6), which harbor key glutenin and gliadin genes.

These observed variations and the presence of contaminant or different alleles at the *Glu-B1*, *Glu-A3*, and *Glu-D3* loci suggest that the ROELFS line with the translocation is not an exact near isogenic line and is still carrying some genes, such as those at the *Glu-D3* locus on the short arm of chromosome 1D, derived likely from the wild relative donor parent harboring the translocation.

However, it is important to mention that the quality of ROELFS-Lr#N-SA was only slightly inferior compared to control. ROELFS is a high-quality variety, and the translocation line is still classified in the CIMMYT grading system as a wheat line with good quality.

Additional backcross steps to ROELFS might be needed as well as studies on their recurrent parent, to better understand the possible linkage between the Lr#N-SA translocation and gluten strength. Further characterization of the grain proteins in the BNI and non-BNI lines using chromatography and proteomic analysis would help to better understand how the protein profile changes in the presence of the translocation.

### Summary and Outlook

The data presented show that BNI can be expressed in alkaline soil which is a very crucial result for the use of BNI in the Global South, where most alkaline soils are located. We suggest that three mechanisms are involved: (I) active release of root exudates with BNI activity triggered by ammonium addition. (II) A passive release during root turnover and (III) an increased nitrate uptake and transport to the flag leaf observed for the translocation line. The latter could be a secondary beneficial effect due to Lr#N-SA. Nitrate reduction after N fertilizer application was measurable in the topsoil and potential nitrification rates (*in vitro*) were clearly linked to *in situ* addition. The hypothesized positive effects of BNI on soil ammonium could not be confirmed. The interpretation of the data remains challenging and calls for studies on ammonium immobilization (net mineralization and nitrification studies) of the nitrifier soil community. The translocation delayed anthesis and maturity. Whether this is a positive or negative alteration by the translocation will need to be assessed in future. Further breeding efforts are necessary to ensure a more consistent grain yield formation and to correct for the lower HI of the translocation line. For the tested line ROELFS-Lr#N-SA (BC_5_), more backcrosses might increase its yield performance and remove the negative allele that causes weak gluten. This study was limited by using only one genotype, ROELFS. It remains crucial to investigate the interaction of the T3BL.3NsbS chromosome-region (Lr#N-SA) trait in other and more recent elite spring wheat backgrounds.

## Author contribution

IO-M, H-JB, MK, AB and HK conceived and designed the experiment. IO-M, HK and MEC-C conducted the experiments. HK, IO-M and MII analyzed the data. VK was responsible for donor relations and edited the manuscript. HK wrote the manuscript. All authors contributed to the discussion of the data set, to the manuscript revision, read, and approved the submitted version.

## Acknowledgements

We are deeply grateful to Tony Fischer for reviewing the manuscript before submission. We acknowledge the contribution of Lorena Velazquez Peredo to this work by conducting the analysis of soil and plant tissue samples. For her collaboration in organizing logistics during the leaf tissue sampling at CENEB we would like to thank Jacinta Gimeno Romeu and Gemma Molero.

